# Understanding melanopsin using bayesian generative models – an Introduction

**DOI:** 10.1101/043273

**Authors:** Benedikt V. Ehinger, Dennis Eickelbeck, Katharina Spoida, Stefan Herlitze, Peter König

## Abstract

Understanding biological processes implies a quantitative description. In recent years a new tool set, Bayesian hierarchical modeling, has seen rapid development. We use these methods to model kinetics of a specific protein in a neuroscience context: melanopsin. Melanopsin is a photoactive protein in retinal ganglion cells. Due to its photoactivity, melanopsin is widely used in optogenetic experiments and an important component in the elucidation of neuronal interactions. Thus it is important to understand the relevant processes and develop mechanistic models. Here, with a focus on methodological aspects, we develop, implement, fit and discuss Bayesian generative models of melanopsin dynamics.

We start with a sketch of a basic model and then translate it into formal probabilistic language. As melanopsin occurs in at least two states, a resting and a firing state, a basic model is defined by a non-stationary two state hidden Markov process. Subsequently we add complexities in the form of (1) a hierarchical extension to fit multiple cells; (2) a wavelength dependency, to investigate the response at different color of light stimulation; (3) an additional third state to investigate whether melanopsin is bi‐ or tri-stable; (4) differences between different sub-types of melanopsin as found in different species. This application of modeling melanopsin dynamics demonstrates several benefits of Bayesian methods. They directly model uncertainty of parameters, are flexible in the distributions and relations of parameters in the modeling, and allow including prior knowledge, for example parameter values based on biochemical data.

## 2. Introduction

Time-varying data can be analyzed with a multitude of statistical methods. Integrating ordinary or partial differential equations is one of the major tools in the natural sciences. For example in order to analyze the morphology of an action potential we could model the rise and fall by a system of two coupled differential equations. In a linear approximation this results in two exponential functions, where the time-constants of the exponential describe the rise and fall. Alternatively we could use the more complex Hodgkin-Huxley model (Hodgkin and Huxley, 1952). This system of equations does not only better describe the data, but allows a direct interpretation of model variables in terms of molecular and cellular properties. Furthermore, in many experiments, multiple factors influence the dependent variable concurrently and the process of interest is non-stationary. In that case, extracting single time constants can be biased and unable to explain the data. And consequently the mechanistic model should be preferred. The benefit of such generative models is the ability to generate ‘fake-data’ using previously fitted parameters. It allows to predict unseen data and simulate experiments where, for example, some of the parameters where changed. Thus, the first step to analyze time-varying data, is to develop a formal mechanistic model of your data.

Once we specified the model, we need to estimate the values for the parameters based on measured data. A solution to such systems of differential equations is most commonly in the form of maximum likelihood estimates, i.e. the one parameter set so that the occurrence of the data as observed is most likely. While often used, another approach has important benefits and improvements: Bayesian parameter estimation. It allows us to directly estimate parameter uncertainties, interpret them intuitively as probabilities about parameters conditioned on the data and we are able to seamlessly include prior knowledge. Due to these benefits, Bayesian parameter estimation has seen a strong comeback and is becoming ever so popular (Cronin et al., 2010; Ghasemi et al., 2011).

In order to use Bayesian estimation we need to understand three concepts: the likelihood, the prior and the posterior. The likelihood tells us how likely it is, that our data are generated by a given set of parameter-values. The prior tells us, how likely certain parameter-values are in the first place. Thus if we *a-priori* know that a receptor has a certain time-constant from previous experiments, we can directly incorporate this knowledge in our current model-fit and adequately influence the posterior of the time-constant parameter and all other co-dependent estimates. The posterior of each parameter is the distribution that shows us how probably each parameter-value is, given our data and prior knowledge, thus a combination of prior and likelihood. In the end we do not only get a single best-fitting parameter value, but a distribution. Thus in addition to the most probable parameter value, we estimate the uncertainty of the parameters, the probability distribution. A broad probability distribution indicates that we cannot estimate the parameter well: neighboring parameter values have a similarly high posterior probability. But a thin distribution indicates that the parameter can be estimate with high precision. Furthermore, dependencies between several parameters might be complex, but can be modelled by these methods. With Bayesian methods we can flexibly use generative models and, importantly, the posterior probability can be interpreted as uncertainty of a parameter, a straight forward and often implicitly used interpretation.

As an example to guide this paper we use patch clamp recordings of cells expressing melanopsin, a photosensitive opsin-type occurring naturally in the retina. In mammals it is expressed in ganglia cells and projects to the suprachiasmatic nucleus and influences the circadian rhythm (Hankins et al., 2008; Do and Yau, 2010). A cell containing melanopsin will begin to fire if photons of a certain wavelength activate the protein. Melanopsin is activated using blue light (470 nm) and can subsequently deactivated using green-yellow light (560 nm). In contrast to other opsins, melanopsins’ activation is tonic, once activated it stays activated for several seconds to minutes (Spoida et al., 2016). Melanopsin presumably occurs in two states, the M (active) and R (resting, inactive) states (for a review see (Schmidt and Kofuji, 2009), but see (Emanuel and Do, 2015)). Activating the protein with blue light increases the probability of the R-state melanopsins to change their configuration to the active M state. Concurrently, a constant transition-probability from R to M and M to R, exists that leads the cell to an equilibrium distribution of melanopsin in M and R state configurations. Here, we use data from melanopsin patch clamp recordings (Spoida et al., 2016), where cells at resting state are activated using blue light and subsequently deactivated with red light (Figure 1)

**Figure 1:**
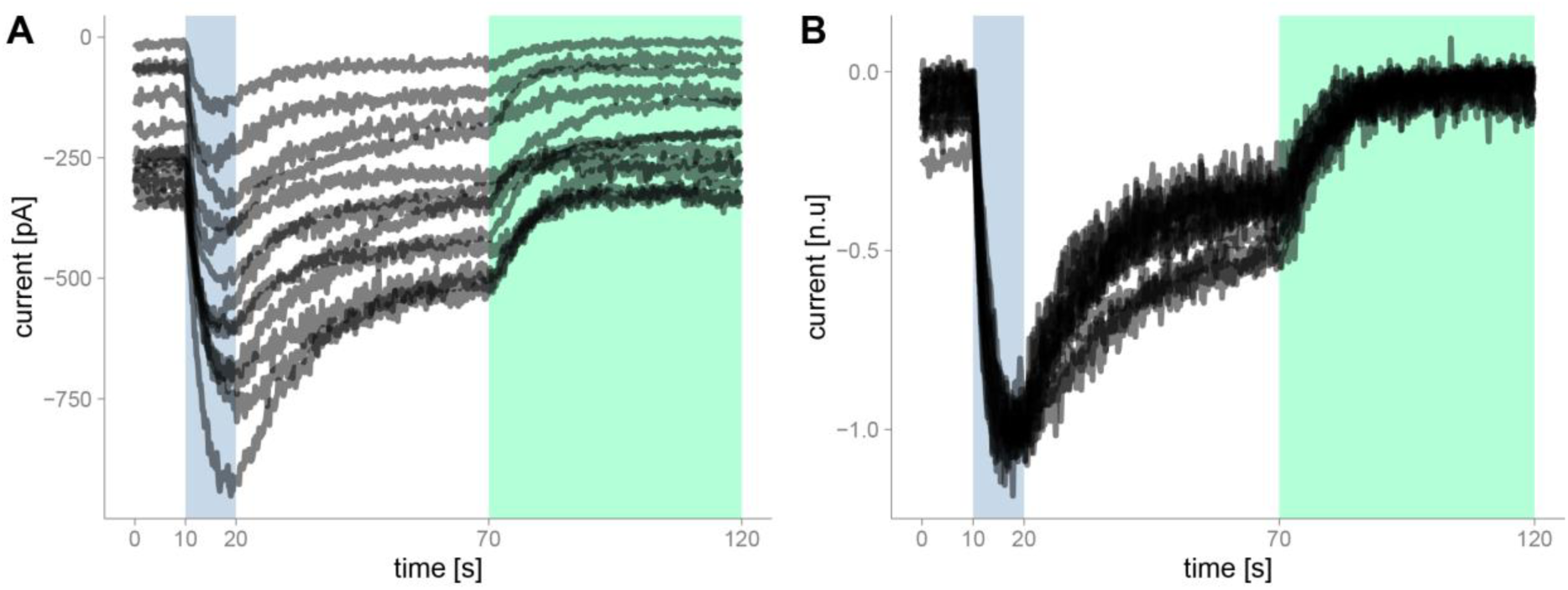
A) Raw data of hOpn4l patch clamp recordings. hOpn4l was expressed in HEK 293 cells which express GIRK1/2 subunits. The GIRK-mediated *K*^+^-currents were sampled at 50-200Hz. Blue light (470nm) activates current outflow, green/yellow light (560nm) deactivates the outflow. B) Data were resampled to 5 Hz. We then normalized the range by mapping the 95% percentile of each cell between 0 and -1.

In this paper we develop a Bayesian mechanistic model of melanopsin and discuss the implementation of the model, the inverse fit, model checks, the interpretation of the parameters and how we can exchange parts of the model in a modular way to improve our understanding and design new experiments.

## 3. Methods and Results

### 1. Model building

It is helpful to start with a graphical model representation (Figure 2 A). In this paper we loosely follow the model notation in (Lee and Wagenmakers, 2014). Once the graphical model is specified, it can be directly implemented into a Bayesian programming language. In the graphical model (Figure 2 A) all parameters that change over time are shown inside the time point – plate and indicated with time-indices. The main parameters are the proportion of firing (M) and resting (R) states. In every simulation time step *t*_*i*_ there is a certain probability to switch states from M to R: *p*(*MtoR*)_*t*_. This transition probability is influenced by a constant rate *C*_*MR*_ and a green-light dependent rate *L*_*MR*_. Of course the light dependent rate is only taken into account, when there is green light, thus we need a dummy-coded green light variable *L*_*G*_ with 0 when there is no light, and 1 when the green light is active. Because light-activation happens at specific times determined by the experimenter, the transition probabilities change over time, i.e. they are non-stationary. The transitions are implemented using ordinary differential equations. One of the assumptions of the model is, that the recorded patch-clamp currents are directly proportional to the proportion of M-state. We don’t expect the patch clamp noise level to change during our recording time, and thus we include a constant Gaussian noise term into our model. To summarize: We model the patch clamp currents using a Gaussian where the mean is proportional on the amount of M-state and thus non-stationary over time. The model allows us to intuitively grasp the parameters, interactions and mechanisms that are needed to model our data.

**Figure 2:**
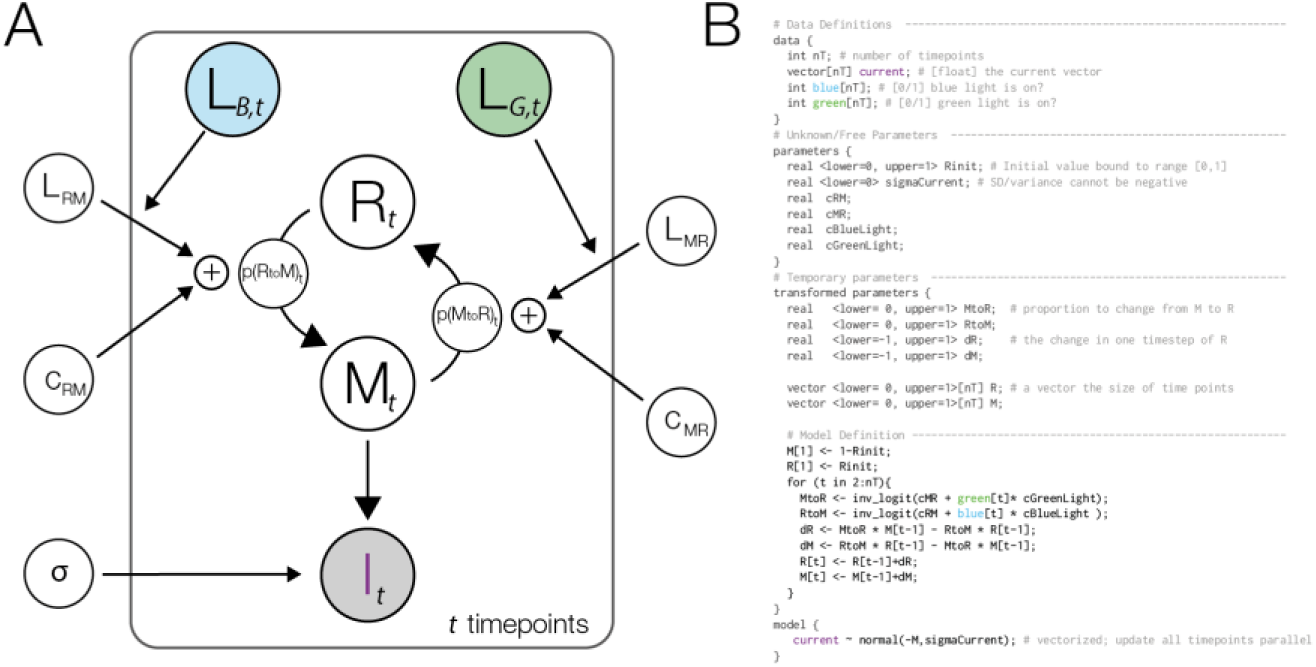
A) A graphical model description of the basic model. Filled parameters depict data that is given. At each point in time a fraction of the R state is changed into the M state with the non-stationary probability p(RtoM). The same process governs the change from M to R. The transition probabilities are influenced by constant (stationary) leakage probabilities and non-stationary, light dependent activations. The active M state is used as a model of the measured current of the patch clamp. These recording are inherently noisy, and we model this noise using a Gaussian function with the non-stationary mean *M*_*t*_ and the standard deviation *σ*. The parameter for the initial M/R state at t=1 was omitted from the graph. B) The graphical model implemented in the STAN programing language.

A more formal way to describe this implementation is to describe the model as a non-stationary two-state hidden Markov model. We then estimate the transition probabilities and relevant factors. All scripts and models are documented and publicly available under http://osf.io/bn6pk. In this paper we make use of the STAN packages (Carpenter et al., 2016), in combination with R (R Core Team, 2013). The non-stationarity in our case was implemented by a logistic linear model with time-varying predictors. The model code is shown in Figure 2 B, parallel to the model graph. In the following, square brackets reflect arrays, round brackets reflect functions. In our case, the linear model can be described by:

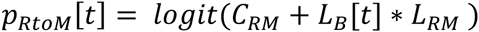

Where *c*_*RM*_ is the constant change parameter, *L*_*B*_[*t*] defines at which time intervals blue light is active and *L*_*RM*_ is the blue light dependent change parameter. The logit function maps values from the domain ‐infinity to infinity to the domain of 0 to 1, thus in the domain of probabilities. This formulation as a logistic linear model allows us to connect the estimation of parameters over multiple cells with the idea of hierarchical or mixed models (see section *Modular Improvements*, hierarchical fit further down). The same formula defines the spontaneous transition probability from M‐ to R-state. Thus for the size of change of R-state at each point in time, there exist two influences: Some fraction of melanopsin changing their state from M to R and in the same time step some spontaneous change from resting state to fire state. The combined probability determines the proportion of R (or M respectively) as captured by using ordinary differential equations. At each simulated time step (with a predefined time-resolution *Δt*) we update our R-parameter (and M respectively) by a first order integration:

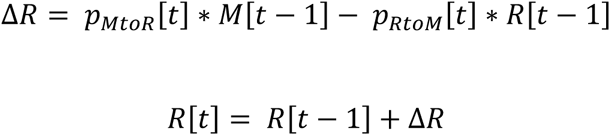

We use a discrete time notation here to parallel the code of the implementation. In the two-state model it is necessary that the amount of M state is equivalent to the inverse of the R state. Thus:

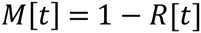

We can make use of this relation and only calculate the change in R state and invert the change in the M state, but if we want to enhance the model to three states, it is more sensible to implement both changes, dR and dM.

The final important relationship to define is the relation to our data and including a noise distribution. In STAN this can be achieved by using:

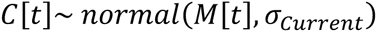

In STAN the tilde (~) means „is sampled from“. Thus, the line defines that the measured current C is sampled from a normal distribution with time-varying mean and constant variance *σ*^2^. Before the model fit we need additional statements about the type and range of parameters, we need to define the initial state, e.g. which could be random.

This concludes the implementation of the specified graphical model into STAN.

### 2. Bayesian Parameter estimation

In the next step we estimate the posterior parameter distributions. Here we will give short introduction of Bayesian data analysis and Monte-Carlo sampling methodology. Our goal is to estimate the posterior probability distribution: colloquially, what is the probability that each possible parameter value could underlie our data. According to the Bayesian framework, this consists of firstly the likelihood of the data given the parameter. In other words how likely is it, that the data are generated from a specific set of parameters. Secondly from the prior distribution which states how probable a parameter is in the first place. In more formal terms, we are interested in the posterior distribution (*p*(*θ*|*D*)) given the likelihood of the data (*p*(*D*|*θ*)) and prior parameter probabilities (*p*(*θ*)). Bayes theorem states that these are directly related to each other (*p*(*θ*|*D*)~*p*(*D*|*θ*) * *p*(*θ*)). An example: We record a neuron spiking with 10Hz. Our imaginative model assumes that the spiking rate of the cell is sampled from a normal distribution with a mean and a fixed standard deviation at 2 Hz. This model has only a single parameter to be estimated. We can easily calculate the likelihood of the Gaussian: We will get a low likelihood for a set of parameters where the mean is 5 Hz, a higher likelihood for a mean of 12 Hz an even higher likelihood for a mean of 10Hz. If we incorporate prior knowledge that these specific neuron types are very rarely observed with a spiking rate of higher than 5Hz, Bayes rule will integrate the information gained from the data and the prior-information and we will find the most likely parameter, given prior and data, at a lower estimate, for example 8 Hz. Whether data or prior dominates the posterior depends on how accurate, or certain, your prior knowledge was specified, and how much uncertainty, or noise, about the parameter the data has. Bayes rule automatically finds the optimal compromise between prior knowledge (what we think is a likely result) and our data (what actually happened).

Calculating the posterior is straight forward for a single parameter: We could randomly try out all parameter values using a grid approach, calculate the likelihoods and priors, and observe the posterior. This would be very ineffective, especially for if we have to estimate multiple parameters concurrently as there is a combinatory explosion. This is where the markov chain monte-carlo (MCMC) sampling comes into play. Instead of randomly sampling the space, we start at a random initial value and propose to jump to a new value. We evaluate the posterior at this value, if it is higher (thus more likely) than the current value, we will go there. If it is lower, we will go there only with a probability inverse proportional to the difference. From there on we repeat the procedure for many iterations. This simple rule (known as the metropolis algorithm, (Hastings, 1970)) will ensure that we visit areas more often where the posterior is high, but from time to time explore other, less probable areas as well. Moreover our Markov chains fulfill all assumptions of the ergodicity theorem, thus it is guaranteed that the Markov chain will *ultimately* converge to the true posterior. In the end, our estimate of the posterior consists of how often we visited a certain parameter value. The MCMC sampling algorithm allows us to estimate highly complex models with many parameters.

Over the years more sophisticated algorithms have been developed. In this paper, we use NUTS, the No-U-Turn sampler, it is more efficient than the metropolis samplers in the case of hierarchical linear models with correlated parameters. This algorithm stems from the family of Hamiltonian monte-carlo (HMCs) algorithms. With HMC algorithms we replace the randomly chosen proposal step of metropolis with an algorithm that more effectively samples the posterior. Imagine that the inverse of the posterior has a bowl shape, thus the most likely points are at the valley, and the most unlikely one raise as mountains the further away you go. We randomly start at a point in the posterior and place a marble and send it with a small push in a random direction on its way. We now simulate for a while and the position the marble ends up, is our new proposed value. We compare it again to the current value and proceed as before. The marble has some momentum so it might just be enough to roll through local minima. The NUTS algorithm is based on HMC but in addition makes certain to not allow any u-turns where the marble rolls uphill (due to gained or initial momentum) and would come down the same way again. The exact algorithm is somewhat more difficult because it needs to make certain that it converges towards the posterior but this is the general idea. For details we refer the interested reader to (Homan and Gelman, 2014). NUTS allows for an effective sampling of the posterior and reduced the risk to get stuck in local minima or passages where the chains could get stuck in the posterior landscape.

### 3. Model Fit & Sampling Diagnostic

Next we describe how STAN estimates the posterior distribution. Stan is a sophisticated open source implementation of HMC/NUTS for a multitude of programing languages (R, Matlab, Python, Julia, Stata and a command line tool). It allows to specify models in a comparatively simple way and has many tools to evaluate the results. The model comes with their own programming language which is not difficult to learn if experience in python, R, matlab or c++ are available. The STAN-model is then compiled to c++ code by the STAN interface and sampled by the MCMC algorithm. Sampling consists of two phases, the first is a warmup period where sampling-parameters are calibrated by the NUTS algorithm to effectively sample from the shape of the posterior. This is necessary as new proposed values could be outside the allowed range of the parameter and in that case we would have to reject this location proposal, thus we have an overhead of likelihood calculations. If this happens too often, we sample ineffectively. But at the same time, we do not want redundancies in the sampling resulting from small (but not rejected) step sizes. This tradeoff is automatically calibrated in the warmup period and the following sampling period defines the final outcome of our posterior.

The chains of an MCMC sampler need to be diagnosed for proper convergence. Sometimes we can get stuck in certain parameter value constellations, for example in a bimodal posterior distribution, or the MCMC algorithm makes too small jumps and we do not explore the space appropriately. It is difficult to diagnose those problems when we only look at a single chain with a single starting value. Therefore we use multiple chains which run independently. This allows us to check whether the chains converged in the same posterior distribution, which is necessary (but not sufficient) for successful sampling. There are several features that can indicate proper convergence: We visually inspect the chains (Figure 3 A), compare the variance between chains to the variance in one chain (termed RHat, and should be close to 1), look at the overlap of the posterior densities of the chains (Figure 3 B) or we check the autocorrelation of a chain (Figure 3 C), how independent two following samples are from each other. A high independency is preferred here. In Stan this is often reported as a single number, the effective number of samples, N_eff, which is the number of samples corrected by the autocorrelogram ((Gelman et al., 2013) p. 286). In our first model, we ran 5 chains with 300 warmup iterations and 500 samples. Visually we see that the chains seem converged and the posterior overlap. Similarly the Rhat is below 1.1 for all parameters. The autocorrelogram shows autocorrelation up to a certain degree, but it does not seem worrying (Figure 3 C, upper panel). In a similar vein, the effective samples are 700 for the upper and 1670 for the lower parameter, representing the ‘best’ and ‘worst’ effective sample value in this model. According to all our criterions, the chains of the MCMC seem to have converged.

**Figure 3:**
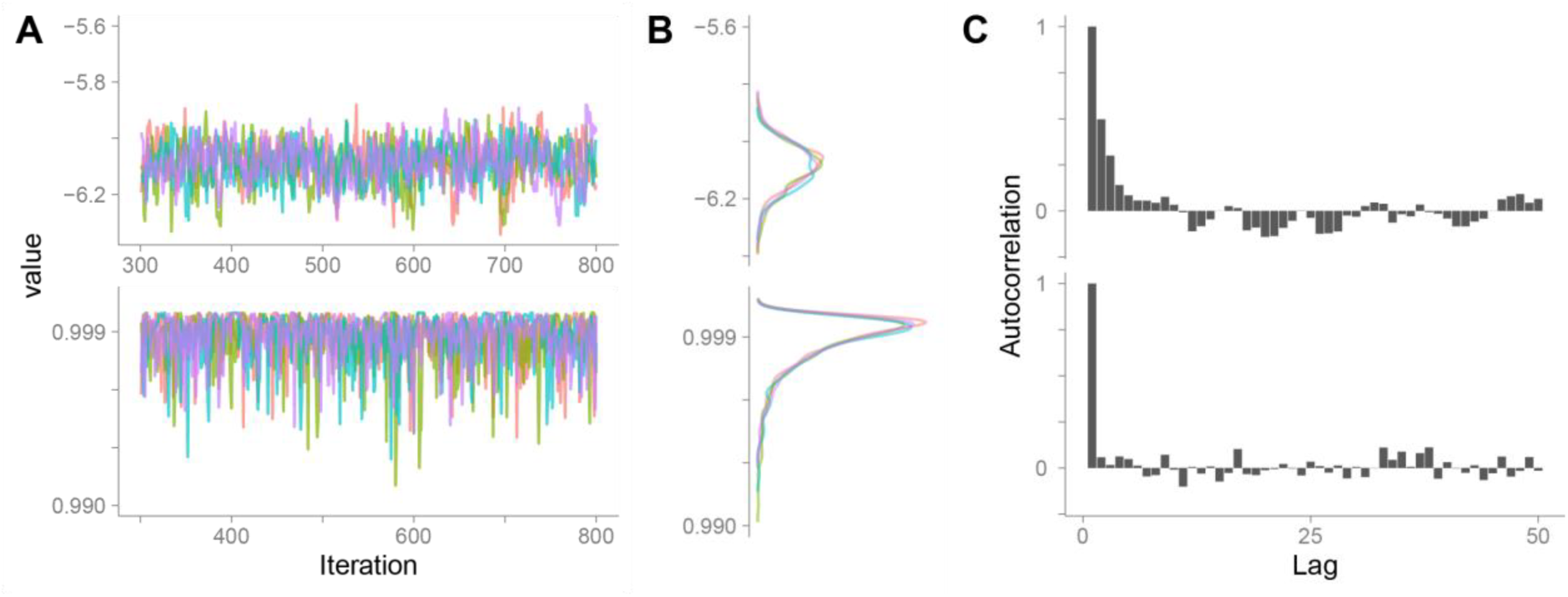
A) four independent MCMC chains with 500 samples each of two parameters, RtoM and Rinit. The chains all converged to the same value range. The variance between chains is similar to the variance within chains. Visually, these chains seem to converge to the same value. B) The posterior density (marginals) of the chains in A. The chains all sample the same region of the posterior, this is an indication for convergence of the chains. These densities can further be simplified by specifying for example the medians and 95% quantiles of the distribution. C) The autocorrelogram of the two parameters. The upper parameter (MtoR) has a higher autocorrelation, thus the effective number of independent samples we drew from the posterior is smaller than for the lower parameter (Rinit).

### 4. Posterior Predictive / Model checks

After we have samples of the posterior distribution and preferably before interpreting the results we need to check the adequacy of our model. A powerful tool of generative models is, that they are able to simulate new data from the current estimated parameters. These new data should capture the important dynamics and effects of our original data. Otherwise the model would be inadequate. When we sample new data from the posterior parameter estimates, this process is called posterior predictive model check. In our example, we sample 1000 new traces from our posterior parameter estimate distributions (Figure 4). Because we randomly sample from a distribution of estimates, each trace will be a little bit different. Our original data should be in the 95% credibility interval of the posterior predictive set. Only after we ensured that the model is adequate, we inspect the posterior parameter estimates visually or calculate and interpret summary statistics (often median and percentiles) of the parameter distributions and interpret the results.

**Figure 4:**
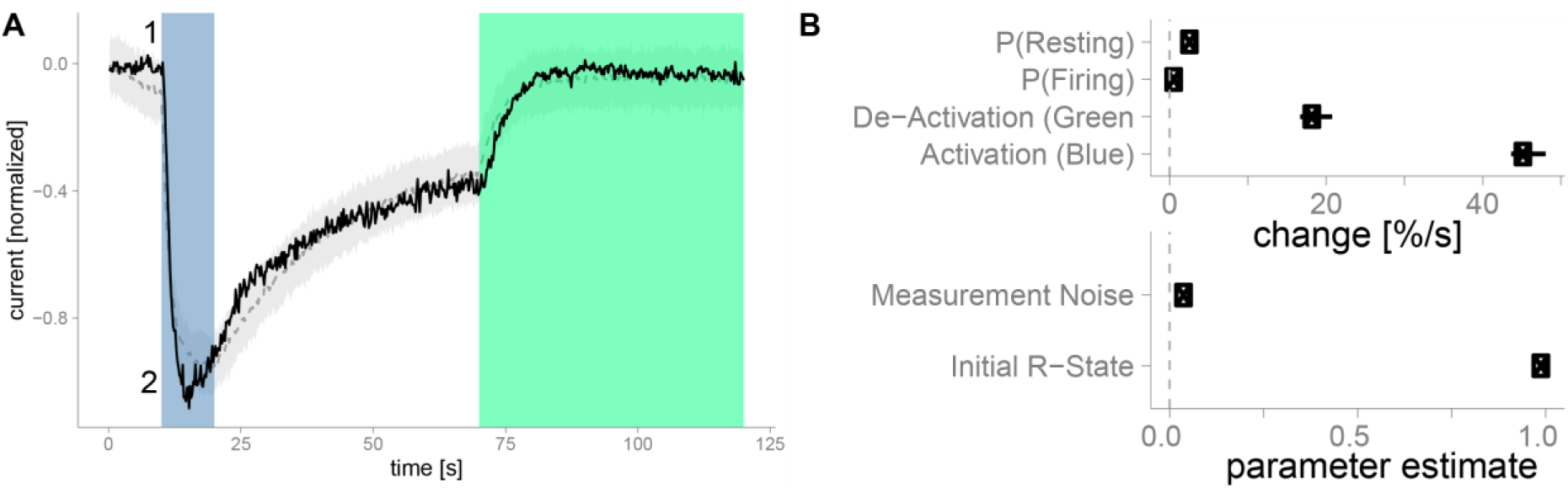
A) The light gray band depicts the 95% credibility interval of 1000 posterior predictives. Posterior predictives are ‘new cells’ that are simulated from our posterior parameter estimates and reflect the range of possible outcomes of the posterior model fit. The dark gray line depicts the median posterior predictive value. The black curve depicts the original data. The annotation “1” and “2” are discussed in the text. B) Parameter estimates of the single cell shown in A). median and 95% percentiles are shown.

The posterior check reveals two problems with our model. In the initial phase, marked with (1), we observe a mismatch between the observed and the predicted data. Here, the posterior predictives indicate that the current is slowly increasing, whereas the data indicate no such trend. This first model missmatch can be readily explained: In the initial phase, the expressed melanopsin proteins are not activated, they need a first activation by blue light, before they can acquire an equilibrium between the R and M state. But the model assumes falsely, that this equilibrium can be acquired from the beginning. By either excluding this portion or adding another initial state for melanopsin, this difference could be modeled. The model mismatch at (2) is currently not well understood. Even though blue light is still activating melanopsin proteins and forcing them to the M state, the current is diminished again. Mechanism that are able to resolve this range from internalization of receptors, to effects of delay due to the g-coupled receptors, due to a hypothetical refractory period of melanopsin or the g-coupled pathway. This cannot be captured by the current model and thus the posterior predictive show an expected maximum at the end of the blue period.

The posterior checks revealed two problems with this model, especially the first one could bias our parameters. To cope with these problems is left open for now, but it is not difficult to resolve them by enhancing the model.

We are now ready to interpret our parameters for this single cell fit. We expected the initial R-state parameter to be around one, due to our baseline correction. This is indeed the case, the average initial state for R is 1 [0.99,1]. The estimated standard deviation (the estimated measurement noise) of our signal is 0.052 [0.050,0.054]. We defined four main parameters in our model: The first is ***c***_***RM***_, it indicates the constant and spontaneous transition probability from the resting to the active state. The estimate is -6.5 [-6.6, -6.5] on the logit scale. In order to convert this to a more sensible unit, first we take the inverse of the logit function. Then we need to raise the 1-x to the sampling frequency to gain the probability per second:

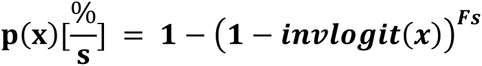

Thus converted to percent per second, the spontaneous change is on average 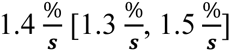. The spontaneous transition back to the resting state ***c***_***MR***_ is the second parameter and for this cell it is a bit higher with on average 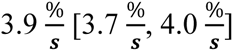. We can also construct the equilibrium point from these data, 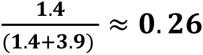, thus we expect the equilibrium state to be at around 26% of the maximal theoretical current (the maximal M state). In order to convert the parameters ***L***_***RM***_, the activation by blue light, one needs to take the concurrent constant change into account, the formula changes to:

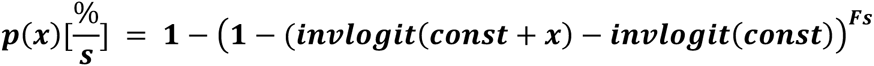

Thus for the activation by blue light we get 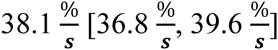 and for green light deactivation we see a change of on average 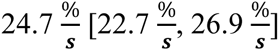. Keep in mind that this is an estimate for a single cell, thus the posteriors are comparably tight, the uncertainty about the parameters is low. More complex models take the data of multiple cells in account and are introduced in the next chapter. This concludes the bayesian model fit. To go further from here we recommend the introduction book by Kruschke (Kruschke, 2014), the applied problem-centered book by Wagenmaker (Lee and Wagenmakers, 2014) and the book by Gelman (Gelman et al., 2013).

## 4. Model Extensions

We are now ready to discuss further enhancements to the model. We advocate to start simple, with a basic working model and after thorough checks, add the modules that are needed for your analysis.

### 1. Hierarchical Model Fit

We successfully estimated parameters for a single cell. Now we need to check whether this hold for the whole population of cells. A standard procedure is estimating the parameters of each cell individually and then taking the average as the population average. This is a valid and straight forward approach, but has some drawbacks: Cells where parameters are difficult to estimate are weighted the same as cells where parameters are certain. In a similar vein, the single cell parameter estimates are not influenced by the parameters of other cells, even though we can leverage this population knowledge to get better single cell estimates. In recent years mixed linear models (also known as hierarchical models) are becoming more and more popular. In mixed models we fit all cells at the same time and assume that the parameter value of each cell is sampled from a parent population parameter-distribution. In Figure 5 A we see that the single cell estimates (green, line shows mean and distribution shows the estimation precision) are samples from an overarching population distribution of the parameter, in this case a normal distribution with two parameters. If all cell-parameters are sampled from the population-distribution, it is reasonable to expect that single cell parameters that are in the tail of the population (thus extreme outcomes or outliers) are unlikely. We thus move our single cell estimate closer to the mean of the population-distribution, an effect termed shrinkage. The amount of shrinkage depends on the probable distribution of the single cell mean and the distance of the cell mean to the population mean (the variance of the population needs to be included in the distance). The population distribution parameters are estimated concurrently to the shrinked single cell estimates. Because we estimated parameters from the same cell, similar to a within-subject design, we expect that multiple parameters could be correlated with each other (Figure 5 C). For example, if we estimate the refractory period of a neuron to be short we might also suspect that it shows a higher maximal firing rate. Thus when estimating multiple parameters of a cell, we have to take correlations between the parameters into account. This also allows for shrinkage over the correlation parameter. If there is no correlation in the data nor prior, the estimate will also be close to zero and shrinkage will not take place. Because population distributions are usually normal distributions we can elegantly assume all parameters are based on a multivariate normal distribution with means, variances and a correlation matrix (or equivalently means and a covariance matrix). This part is equivalent to a linear mixed model where all parameters have random slopes and the complete correlation matrix is estimated.

**Figure 5:**
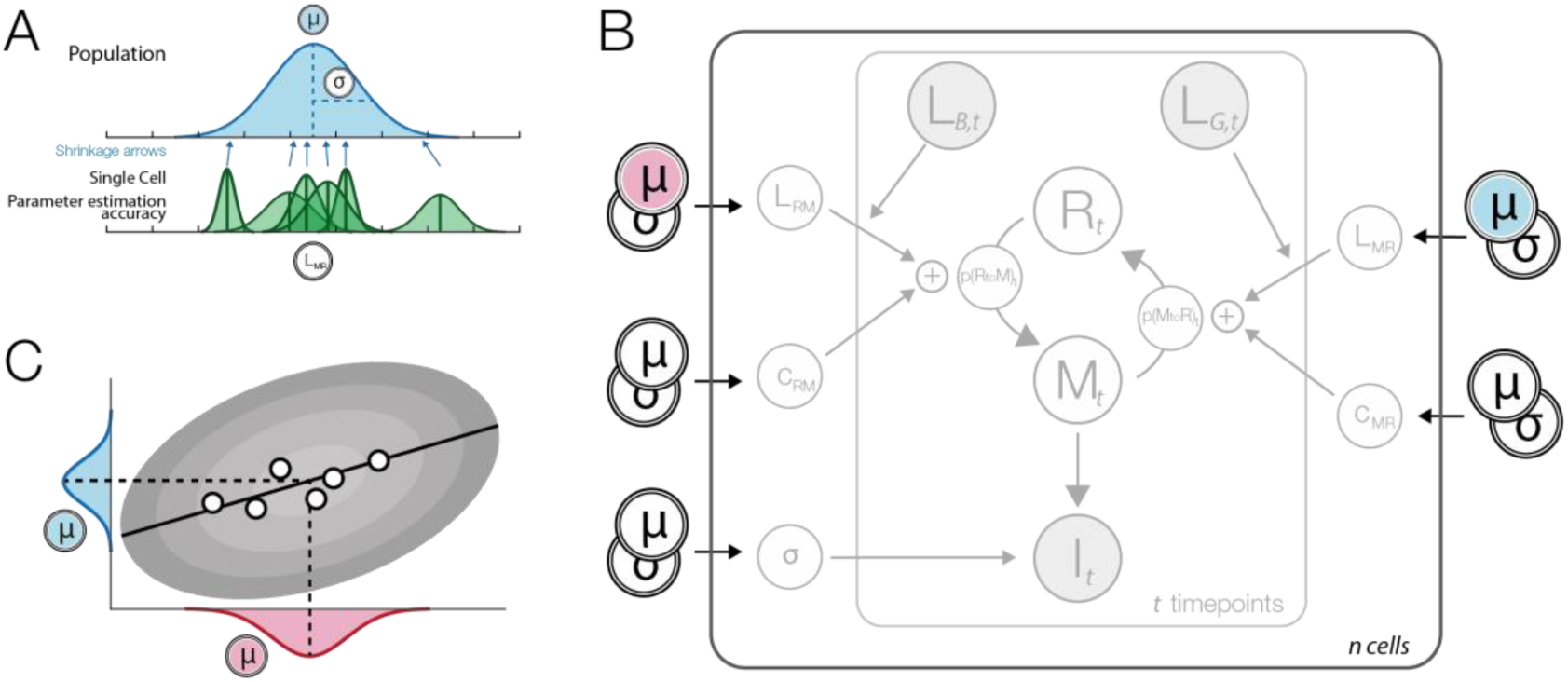
A) Hierarchical parameters. The blue distribution is our population distribution with two parameters, ***μ*** and ***σ***. Below the posterior estimates of the single cells are shown in green. They give an estimate of the mean and its uncertainty. When fitting a hierarchical model, the single cell posterior means are shrunk towards the population mean (blue arrows). Shrinkage is strongest for uncertain (broad distribution) parameters and parameters that are furthest away from the population mean. B) Hierarchical model graph. The parameters of a single cell (see Figure 2) are assumed to be sampled from an overarching population distribution. Thus each single-cell parameter is assumed to come from a population with mean and variance as shown in A). C) Dots represent parameter estimates of single cells. Here we observe a correlation between two population parameters. In order to capture this relationship, we need to include the correlation term between all population parameters and model it as one combined multivariate normal distribution

In practical terms we need to introduce some more parameters to be estimated. The model is kept untouched for the critical calculations in each time step, but of course the underlying data and the parameters are different for each cell. We introduce a matrix notation in the code, where the parameters are saved in a matrix termed beta (dimensions n-parameter * n-cells). We also introduce population-vectors with the prefix ‘m_’ or ‘s_’ for population mean value or population standard deviation value, for example m_beta with n-parameters for the mean. Further we need a correlation matrix (n-parameter * n-parameter) and a population-vector for the variance (dimensions n-parameter, needs to be positive). In stan we can conveniently calculate the covariance matrix using:

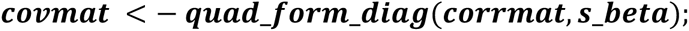

Finally we need to define the relation of the single cell parameters with the multivariate population:

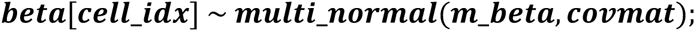

This statement is repeated for each cell through a loop. This states that the beta values (the n-parameter dimension is vectorized, thus hidden) are sampled from a multivariate normal with the given mean and covariance matrix.

The initial value of R for each cell has to be between 0 and 1. But if we sample from a normal distribution with mean 0.9 and SD of 0.1, we will sometimes sample values greater than 0. We can simply ignore those values and in those cases resample until we get a value <1. Alternatively we can use a function that is strictly bounded between 0 and 1, for example a beta-distribution:

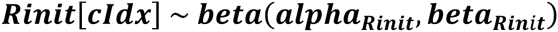

Using pairwise scatter plots of the MCMC values, we noticed that two parameters of the posterior estimates are highly correlated: ***c***_***RM***_, the spontaneous firing rate, and ***L***_***RM***_ the activation through blue light. The correlation stems from the linear model definition and due to the logit scale. In order to activate the cell by blue light, ***L***_***RM***_ needs to act against the very large negative number of ***c***_***RM***_ (a large negative number on the logit scale forces the constant firing probability of the cell to be close to 0). The change at each point in time is:

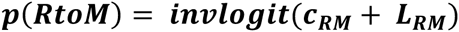

Thus ***L***_***RM***_ needs to counteract ***c***_***RM***_. Let’s take for example 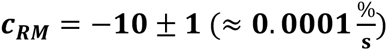.

When light activates the cell, the total should be around 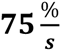:

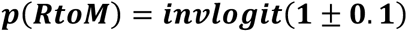

Therefore it is clear that ***L***_***RM***_ **= 11 ± 1.1**. Because the value of ***c***_***RM***_ is expectedly very negative on the logit scale, it will always be a large part of ***L***_***RM***_ and therefore we get the correlations. This is problematic for MCMC sampling algorithms, they do not converge well with high correlations between parameters. There is a trick to reduce the correlation: reparameterization. Reparameterization changes how parameters are related to each other. It only changes the sampling procedure, but not the outcome or the estimated model because we keep the relation between parameters the same. In this case we change:

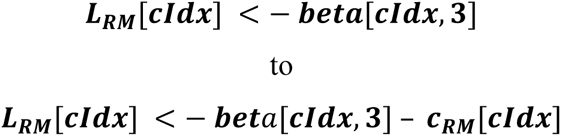

Because we sample beta and not ***L***_***RM***_, we changed the parameter-space that is sampled by the MCMC algorithm, to one that does not show the high correlation between parameters, but we don’t change the actual parameter value. The reparameterization greatly reduced the time to convergence and in addition improved the effective samples ***N***_***eff***_.

With some simple addition to the model we are now able to estimate shrinked parameter values for all cells concurrently. This model is more complex than the simple model, in order for it to converge we needed to initialize the chains at values in the range of the posterior, we used the same values on both the single cell and the population level and initialized the means but not the variances.

It is now necessary to draw posterior predictives to evaluate whether our model is adequate. In hierarchical models, we can perform posterior predictives in at least two cases: either we take the estimated parameters of each cell and do the same procedure as in the basic model for each cell, or we sample “new cells” from the estimated population multivariate normal distribution. These predicted new cells reflect the range of possible results predicted by our model, prior and parameter estimates. For ease of display, we directly plot the amount of M state without the additional noise term added. The first case, selecting the parameters of the single cell, can be seen in Figure 6A. Here the posterior predictives match the real data (Figure 1 B) very well. In the second case we sample new cells, as expected, this results in a broader distribution (Figure 6 B). The general shape again matches the original data very closely.

**Figure 6:**
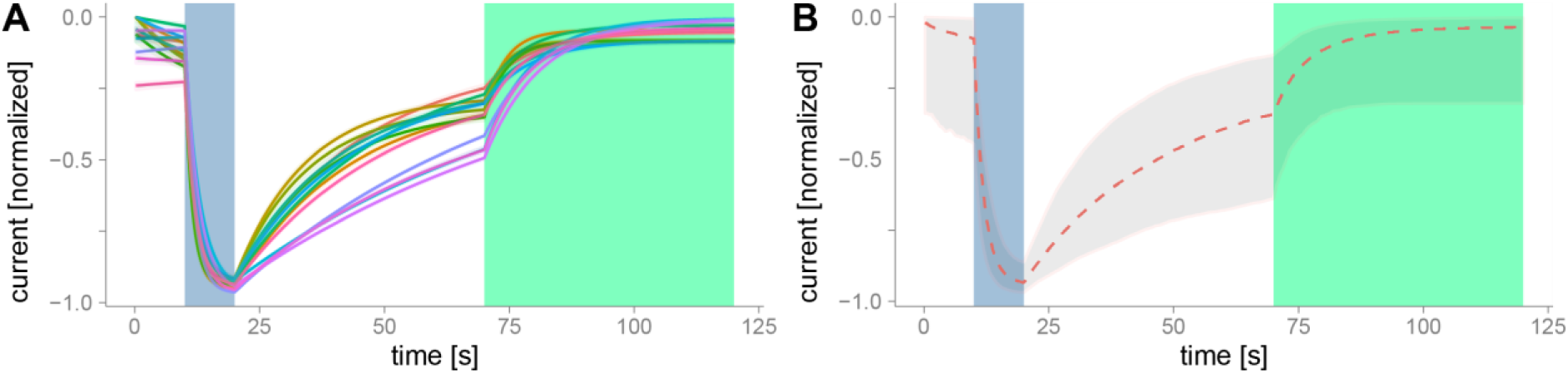
The shaded region depicts the 95% interval of posterior predictives A) Single cell posterior predictive. The parameter estimate of each cell was used to sample new timeseries. Posterior predictives are very similar for single cells, thus the shaded region is nearly invisible. B) Posterior predictive if we sample new cells from the population distribution. Compare with Figure 1 to see the similarity.

After the model posterior predictive tests we look at the results of the model. Similarly to the posterior predictive we can observe results at two different levels. Those two levels, single cell and population, can be seen in figure Figure 7 A,B. In the top posterior estimates of the population distribution, the median distribution and the mean +-95% credibility interval of the mean are shown. In the lower row the single cell uncertainty estimates and their respective means are shown. The population distribution should match the distribution of the single cells, as is the case in A, for ***L***_***MR***_ and in B for the beta-distribution of the initial R-state.

**Figure 7:**
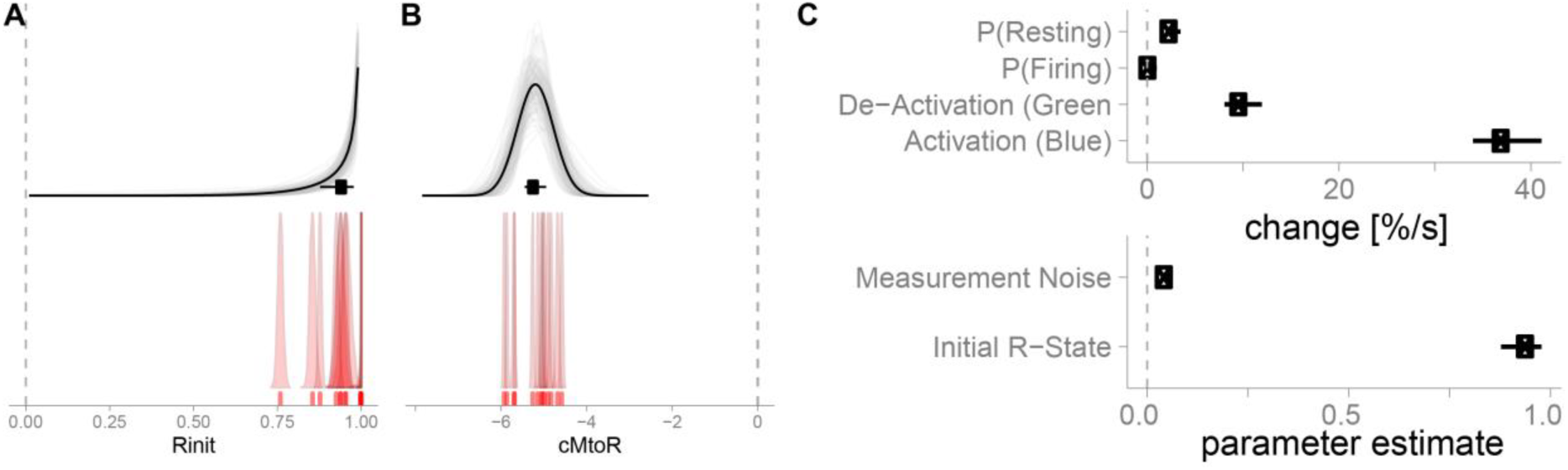
A,B) Population distribution of deactivation with green light (***L***_***MR***_) and the initial R state (***R***_***init***_) respectively with 100 redraws from the posterior chains, the pointrange depicts the mean and 95%-percentile. The lower plots depict single cell posterior estimates and respecte mean posterior. ***L***_***MR***_ is depicted on the logit-scale. C) Results of all parameters. The top plot is in 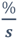, the lower in natural units for the respective parameters.

We can summarize the values using median and 95% percentiles as in Figure 7 C. The spontaneous firing rate is 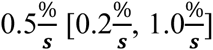, while the spontaneous change to the resting state is 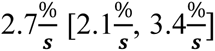, thus the equilibrium point of the population is at 17.4% [8.6%, 28.6%]. Activation by blue light changes the transition probability by 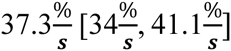. whereas green light deactivates with a lesser rate of by 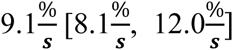. We can also estimate the probabilities of the cell we fitted in the beginning, which will be affected by the shrinkage factor. Here we see that the single cell estimate of the spontaneous firing rate was 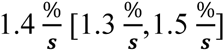 but in the hierarchical model it is 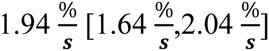. Thus the shrinkage moved the single cell estimate towards the population-mean of 2.7%. This new estimate will be a better prediction of a new measurement of the same cell because it is informed by the estimates of all other cells via shrinkage.

In order to fit multiple cells we needed to add hierarchical population distributions and use a reparameterization-trick. From the model we can sample new cells and estimate in what range new cells will be.

### 2. Priors

Another strength of Bayesian data analysis is the possibility to add prior knowledge to your data. In STAN this is straight forward, for example if we expect that our estimated noise-level is around 0.02 with a standard deviation of 0.01 we add in the STAN-model block:

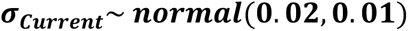

The MCMC sampler incorporates this prior in the appropriate way and integrates it with the likelihood of the standard deviation of the data. Importantly, if we would use a uniform-prior, for example 0.01 – 0.03, we restrict the domain of possible parameter values. Thus even if we have strong evidence from the data that the standard deviation should be 0.05, our posterior will not be able to put any weight, because the prior is zero. This cannot happen with the above normal distribution, because the normal has non-zero weight (albeit very small) from minus infinity to infinity. Another more elaborate example could be to include previously measured biological constants into the model. For example Emanuel and Do 2015 (Emanuel and Do, 2015) proposed a numerical three state model for melanopsin based on biochemical data. They make used of photon absorption rates, spectral templates and quantum efficiencies to simulate the wavelength dependencies of the distribution of states. They then qualitatively compared it to their data and concluded that melanopsin can occur in three states. It is very well possible to enhance the model and include these biochemical data as priors in the data fit and estimate the certainty of the posterior. Priors allow to appropriately incorporate scientific knowledge already at the stage of data fitting.

### 3. Wavelength Dependencies

So far we activated and deactivated melanopsin using two distinct wavelengths. But we can repeat this process with many other wavelengths as well. In that case we are interested to model an activation and a deactivation function of melanopsin based on the wavelength. Of course this function is a priori unknown. While it is possible to use non-parametric basis-functions (e.g. splines) to estimate a non-linear form of the function, in our case there is reasonable evidence (Emanuel and Do, 2015; Spoida et al., 2016), that the activation function follows a Gaussian tuning function. We incorporate this in our model (Figure 8 A) and decided to use a Gaussian with three unknown parameters: a mean, a variance (those two parameters regulate at which wavelengths the cell get de/activated) and a normalization parameter which regulates the strength of the de/activation (Figure 8 B).

**Figure 8:**
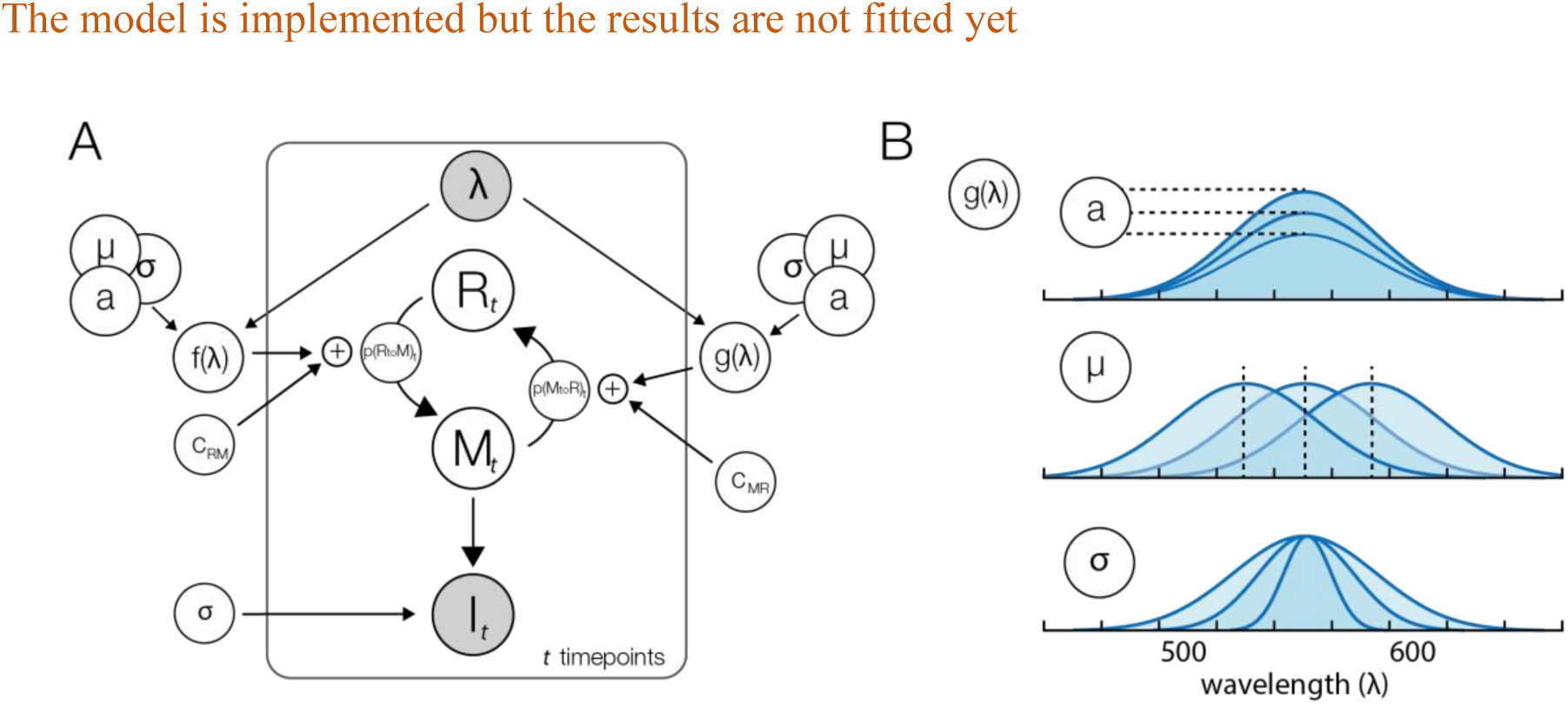
A) Graphical model with wavelength dependency. We replaced the two light sources with a single one that is able to change the wavelength and two functions that translate the wavelength to an activation or deactivation probability. B) The wavelength functions have three parameters. Parameter *a* regulates how strongly the de/activation is. Parameter ***μ*** regulates at which location the maximal de/activation is to be expected and parameter ***σ*** regulates on what range the de/activation can occur.

### 4. Bi‐ vs tri-stability

It has recently be suggested, that Melanopsin has not two states but a third one (Emanuel and Do, 2015). In that case parts of the M state transfiguration change not to the R state, but to the E (extramelanopsin) configuration. In analogue to Emanuel & Do who proposed a numerical three state model simulation, we add this third state X. Therefore the model changes as follows:

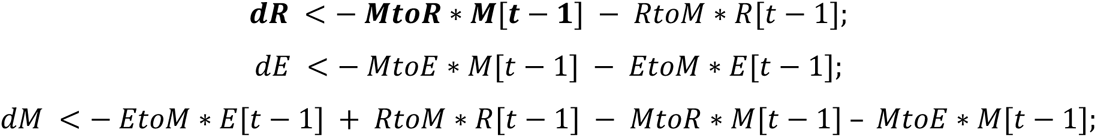

In this model specification transitions from R to E and vice versa are not allowed, which is grounded in energetic constraints where a direct conversion from R to E state has not been observed (Matsuyama et al., 2012). We could perform bayesian model selection on the two state against the three state model to see which model shows more support from the data. This can generally be done using the bayes-factor or an information criterion for example DIC or WAIC. A discussion of the differences or preferences can be found for example in (Gelman et al., 2013). It is to be expected that we need similar data as (Emanuel and Do, 2015) to be able to show that melanopsin has indeed three state. If our current data is already well explained by two states, adding a third state will not improve the model-fit, if we punished for using the additional number of parameters. Indeed a model with three states of a single cell has a WAIC of -5135, while the two state models has only -3696, where a higher number is better. This does not indicate that melanopsin has two states, only that two states are adequate to describe the very limited data gained from a single cell. This module shows the extension of our basic model to be able to directly test two competing hypothesis in a single coherent framework of data analysis.

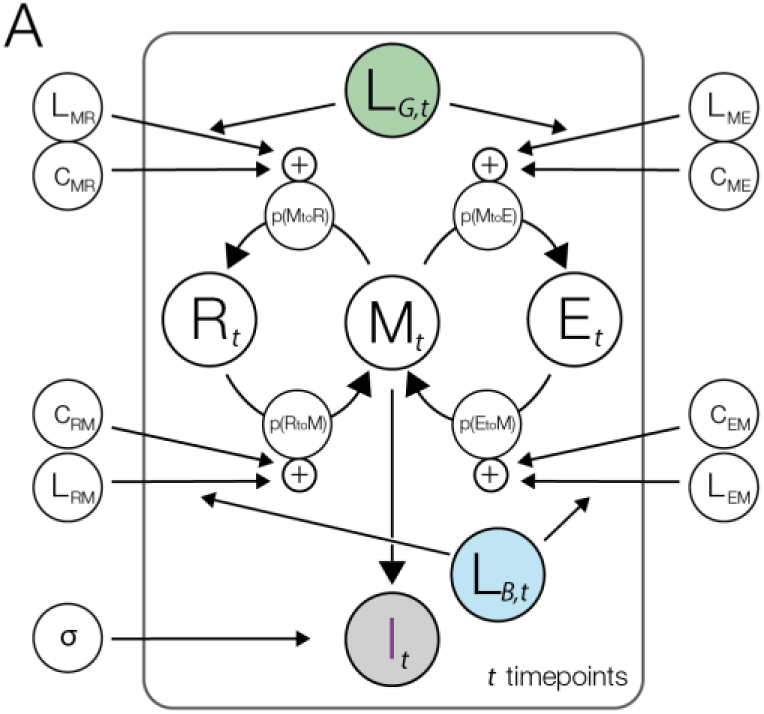

### 5. Differences between cell types

Melanopsin occurs in different species and has slightly different sequences. Two types can show different activation dynamics and thus different underlying kinetic parameters. We recorded data that allows us to compare a human melanopsin (*hOpn4l*) to a mouse-origin melanopsin (*mOpn4l*). We use the basic model with the hierarchical model extension for multiple cells. In our model, we can include this as a factor in the linear model. Thus we adapt two lines in our code:

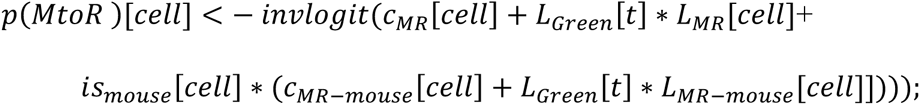

And the respective p(RtoM) line as well. The idea is to model both cell-types with the same parameters, but allow the parameters to differ if the data of a *mOpn4l* cell is being fitted. This is the same way one would model this with treatment coding in a classic linear model. We end up with the parameters for a *hOpn4l* cell (*c*_*MR*_ and *L*_*MR*_) and the difference in the parameters to a *mOpn4l* cell *c*_*MR–mouse*_ and *L*_*MR–mouse*_). If we would like get the parameter estimate for *mOpn4l* directly, we can simply add the two estimates. The results of this model can be seen in the Figure 9 B.

**Figure 9:**
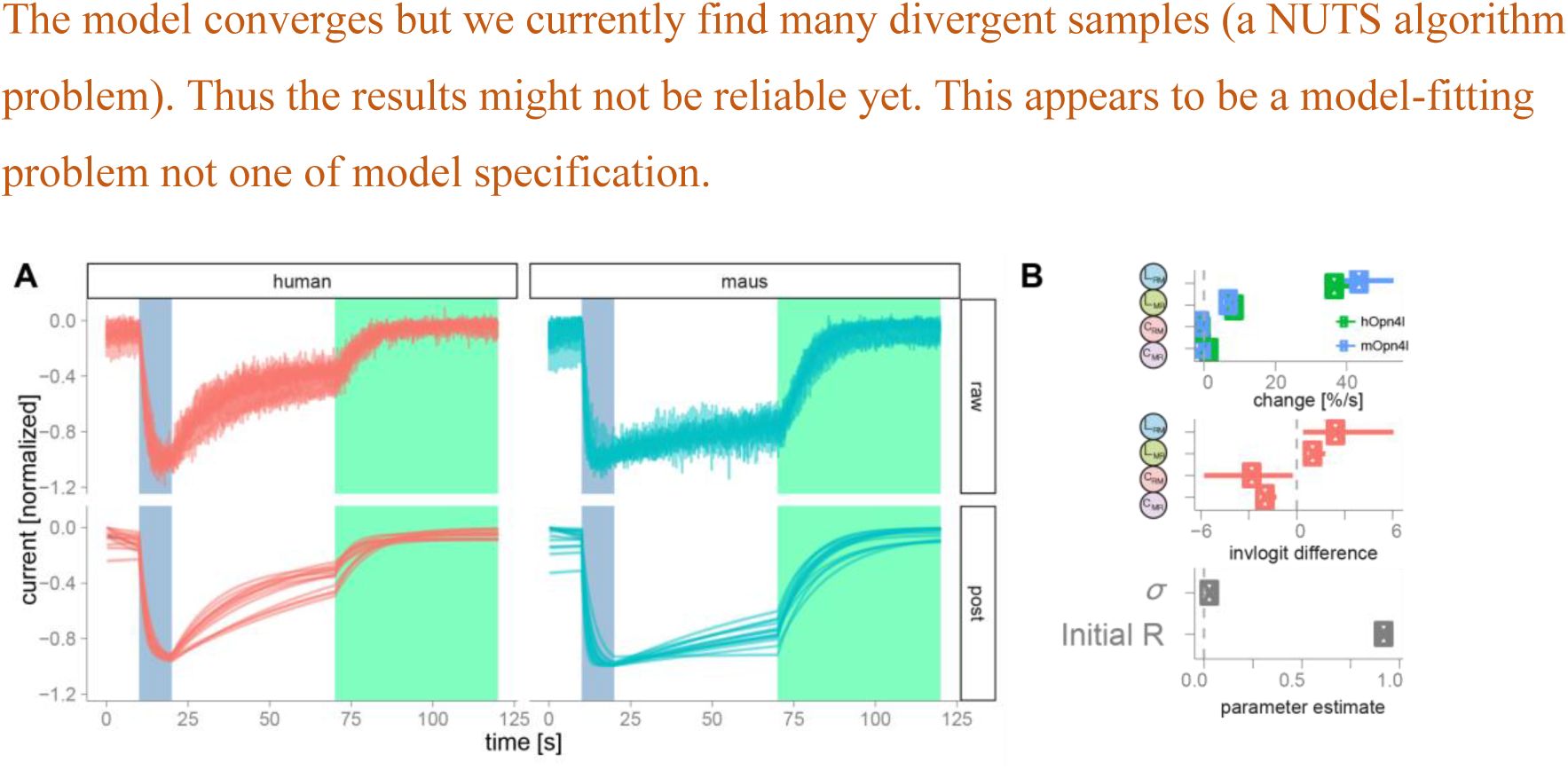
A) The red curves depict 13 cells with human melanopsin (*hOpn4l*). The blue curves depict 14 mouse melanopsin (*mOpn4l*). The upper panel depicts the preprocessed raw data, the lower panel the mean posterior predictive model fit. B) The plot depicts the parameter and their differences of a combined model fit of *hOpn4l* and *mOpn4l*

## 5. Conclusions

In this paper we developed a basic generative model for the kinetics of de‐ and activation of melanopsin. We inverted the model using bayesian parameter estimation in the STAN framework and show how to interpret the parameters of the model and how to predict future data from the model. Using our generative model we are able to inform new experiments and directly tackle uncertainties of underlying parameters.

## References

Carpenter B, Gelman A, Hoffman M, Lee D, Goodrich B, Betancourt M, Brubaker MA, Guo J, Li P, Riddell A (2016) Stan: A probabilistic programming language. J Stat Softw.

Cronin B, Stevenson I, Sur M, Körding KP (2010) Hierarchical Bayesian modeling and Markov chain Monte Carlo sampling for tuning-curve analysis. J Neurophysiol.

Do MTH, Yau K-W (2010) Intrinsically photosensitive retinal ganglion cells. Physiol Rev 90:1547–1581.

Emanuel AJ, Do MTH (2015) Melanopsin Tristability for Sustained and Broadband Article Melanopsin Tristability for Sustained and Broadband Phototransduction. Neuron 85:1043–1055.

Gelman A, Carlin JB, Stern HS, Dunson DB, Vehtari A, Rubin DB (2013) Bayesian Data Analysis, Third Edition.

Ghasemi O, Lindsey ML, Yang T, Nguyen N, Huang Y, Jin Y-F (2011) Bayesian parameter estimation for nonlinear modelling of biological pathways. BMC Syst Biol 5 Suppl 3:S9.

Hankins M, Peirson S, Foster R (2008) Melanopsin: an exciting photopigment. Trends Neurosci.

Hastings WK (1970) Monte Carlo sampling methods using Markov chains and their applications. Biometrika 57:97–109.

Hodgkin AL, Huxley AF (1952) A quantitative description of membrane current and its application to conduction and excitation in nerve. J Physiol 117:500–544.

Homan M, Gelman A (2014) The no-U-turn sampler: Adaptively setting path lengths in Hamiltonian Monte Carlo. J Mach Learn Res.

Kruschke J (2014) Doing Bayesian Data Analysis: A Tutorial with R, JAGS, and Stan.

Lee MD, Wagenmakers E-J (2014) Bayesian Cognitive Modeling: A Practical Course.

Matsuyama T, Yamashita T, Imamoto Y, Shichida Y (2012) Photochemical properties of mammalian melanopsin. Biochemistry 51:5454–5462.

R Core Team (2013) R: A Language and Environment for Statistical Computing.

Schmidt TM, Kofuji P (2009) Functional and morphological differences among intrinsically photosensitive retinal ganglion cells. J Neurosci 29:476–482.

Spoida K, Eickelbeck D, Karapinar R, Eckardt T, Jancke D, Ehinger B, König P, Dalkara D, Herlitze S, Masseck OA (2016) Melanopsin variants as intrinsic optogenetic on and off switches for transient versus sustained activation of G protein pathways. Curr Biol.

